# A mechanism of melanogenesis mediated by E-cadherin downregulation and its involvement in solar lentigines

**DOI:** 10.1101/2023.01.09.523359

**Authors:** Daigo Inoue, Tomomi Narita, Keiko Ishikawa, Katsuyuki Maeno, Akira Motoyama, Takayuki Ono, Hirofumi Aoki, Takako Shibata

## Abstract

**Background:** Intensive studies have revealed pleiotropic melanocytic factors for age spot formation. In particular, dysfunctional keratinocyte differentiation is thought to be an upstream cause of age spot formation. Although keratinocyte differentiation is mediated by a cell-cell contact factor, E-cadherin, its involvement in age spots remains unknown. To find the origin of age spots and an integrated solution, we focused on E-cadherin.

**Methods:** Immunofluorescent staining with cutaneous tissues and cultured cells was performed. Keratinocytes treated with siRNAs were cocultured with melanocytes. With the supernatants of the keratinocyte culture, secretion factors were identified using proteomic analysis. For the activity of melanogenesis and the ingredient screening, a quantitative PCR was performed. For the behavioral analysis of melanocytes, time-lapse imaging of melanocytes was done by confocal laser scanning microscopy.

**Results:** In age spots, E-cadherin expression in the epidermis was downregulated, suggesting that E-cadherin is implicated in age spot formation. E-cadherin knockdown (E-cad-KD) keratinocytes not only promoted the secretion of melanocytic/inflammatory factors, but also increased melanogenesis by upregulating the expression of melanogenesis factors. Furthermore, live imaging showed E-cadherin downregulation detained melanocyte dynamics and accelerated melanin-uptake. Finally, we identified Rosa multiflora fruit extract as a solution for upregulating E-cadherin in keratinocytes.

**Conclusion:** Our findings showed that E-cadherin downregulation triggers various downstream melanocytic processes such as secretion of melanocytic factors and melanogenesis. Additionally, we showed that Rosa multiflora fruit extract upregulates E-cadherin expression in keratinocytes.

## Introduction

Solar lentigo (hereafter called age spots) is a characteristic feature of photo-aging. So far pleiotropic melanocytic environment has been thought to be involved in age spots from the epidermis and dermis. Particularly, in the epidermis, various histological alterations occur such as hyperpigmented basal keratinocytes, rete ridge, dysfunction of keratinocyte differentiation, and possible over-proliferation of keratinocytes [1][2]. These resultant or causal features of age spots are mainly caused by the cumulative effects of chronic UVA and UVB exposure over a lifetime. Since UVB mainly affects the epidermis, mechanistic studies have focused intensively on melanogenesis and its stimulating-signaling in either melanocytes or keratinocytes upon UVB exposure. Upon UVB exposure, keratinocytes promote secreting various cytokines, growth/hormonal factors such as IL-1α, Endothelin-1, and α-MSH [3]. Melanocytic factors bind as ligands to the receptors of melanocytes such as MC1R and c-Kit to stimulate downstream signaling cascades for melanogenesis [4][5]. In melanocytes, MAPK or cAMP signaling cascades activate the transcription factor MITF to promote the expression of essential factors such as TRP1, Tyrosinase, and Pmel17, thereby synthesizing melanosome formation [4][5]. After the maturation of melanosome, melanocytes transfer melanin [6]. Keratinocytes in turn ingest melanosomes and store melanin to protect the cells from UVB-mediated DNA damage [7]. Previous studies have revealed the general mechanism of pigmentation in sun-exposed skin, which also shares a common molecular fingerprint with age spots [8][9][10]. Therefore, it is crucial to investigate how the molecular fingerprint of age spots is different from that of the perilesional area.

So far, the mechanistic overview of the formation and maintenance of age spots consists of four processes: (1) Failure of keratinocyte differentiation [11][12], (2) Upregulation and secretion of various inflammatory, hormonal, and growth factors [6][13][14], (3) Promotion of melanogenesis in melanocytes by the above keratinocyte-derived factors [15][16], (4) Degradation of the basement membrane due to inflammatory factors such as MMP-1, MMP-9, Heparanase [17][18]. Among those, processes (2) and (3) share common molecular factors with sun-exposed non-lesion area but are constitutively modulated for the formation of age spots. Specifically, chronic microinflammation is thought to contribute to forming the melanocytic environment by hindering the differentiation of keratinocytes [19][20]. In turn, the secreted melanocytic factors constitutively stimulate melanocytes. Concomitant with this, proliferative basal keratinocytes excessively store melanin, which arrests the cell cycle of keratinocytes [21]. As a result, abnormal cornification occurs in age spots due to the stagnation of keratinocyte turnover [11][12]. Furthermore, our previous studies have also revealed mechanistic insights into process (4) such as the destruction of the basement membrane and the abnormal formation of a microvascular structure in the dermis [22][23]. Since keratinocyte-derived inflammatory factors have also been suggested to contribute to those abnormalities, keratinocytes are like the point of origin for the creation of the melanocytic environment of age spots. Although these four processes of age spots could directly or indirectly link with each other, how abnormal keratinocyte differentiation in process (1) occurs and is maintained has still largely remained unknown. Recently, one of the key factors for keratinocyte differentiation, autophagy was shown to be critical for melanin degradation, which could lead to the stagnant deposition of melanin in age spots [24][25]. However, the causal factor for dysfunctional keratinocyte differentiation in age spots has remained elusive. Thus, to improve age spots, an integrated solution targeting the origin of age spots has long been sought out.

Differentiation of keratinocytes is regulated by the Notch pathway and the surge of extracellular calcium [26][27]. Upregulation of both the Notch pathway and calcium further promotes transcription for differentiation factors of keratinocytes [26][27]. At the same time, cell adhesion molecules such as E-cadherin, P-cadherin, and Desmocollin-1 are important regulators for keratinocyte differentiation by modulating cell-cell interactions such as keratinocyte-keratinocyte and keratinocyte-melanocyte interactions [28][29][30]. Furthermore, E-cadherin is not only critical for the differentiation/proliferation states of keratinocytes and melanocytes, but also for the survival of the epidermal cells [31]. So far, downregulation of Notch-1 in age spots has been indicated to be an upstream factor for dysfunctional differentiation, indirectly forming a downstream melanocytic environment [12]. However, whether other factors crucial for differentiation are also modulated in age spots remains to be addressed. Thus, to find the origin of age spots and an integrated solution, we focused on a critical regulator of keratinocyte differentiation, E-cadherin, revealing a novel mechanism and solution for improving age spots.

## Materials and Methods

### Assessment of solar lentigines of the human cutaneous specimens

Cutaneous specimens were obtained from the backs of 8 volunteers (Japanese males, 31–49 years of age, average age; 43 years old) with the approval of the ethical committee of the Shiseido Research Center. The diagnosis of solar lentigo was made by a dermatology specialist with the aid of dermoscopy, and 3-mm punch biopsy specimens that included the epidermis with the upper dermis were obtained under anesthesia. Two pairs of specimens were obtained from solar lentigo on the back and neighboring normal sun-exposed skin from individual volunteers. Specimens were embedded into an OCT compound (Sakura Finetechnical, Tokyo, Japan) for the preparation of histological sections. Commercial cutaneous specimens of faces (Obio, LLC. and CTIBiotech) containing hyperpigmented spots were also used by assessing the histological features of age spots such as rete ridge, hyperpigmentation in basal keratinocytes, and thickened epidermis. Among those, the selected samples (N=5, two males and three females, 79–91 years of age, average age; 85 years old)) were embedded in paraffin and subjected to fluorescent immunostaining to examine the expression patterns of E-cadherin and other proteins as in the diagnosed age spot samples.

### Cell cultures

Normal human epidermal keratinocytes (Kurabo) from Caucasian neonatal foreskin were cultured with EpiLife medium (Thermo Fisher Scientific) supplemented with 60 µM CaCl2 and HuMedia KG growth factor kit (Kurabo). Normal human epidermal melanocytes (Thermo Fisher Scientific) from a lightly pigmented neonatal donor were cultured with Medium254 (Thermo Fisher Scientific) supplemented with HMGS2 growth-factor kit (Kurabo). Keratinocytes and melanocytes were cocultured with CnT-PRIME KM (CELLnTEC) supplemented with 1mM CaCl_2_.

### Inhibition of E-cadherin function by E-cadherin neutralizing antibody

For the inhibition of E-cadherin in either cultured keratinocytes or cultured melanocytes, a neutralizing antibody against E-cadherin (SHE78-7, Calbiochem) was used at the final concentration of either 1 or 3 µm/ml. As a control, mouse IgG2a (02-6200, Thermo Fisher Scientific) was used. Either keratinocytes or melanocytes were incubated in the coculture medium supplemented with 1 mM CaCl_2_ for two days, followed by the treatment with either the neutralizing or control antibody for another two days (keratinocytes) or 7 days (melanocytes). For the viable cell counting, fluorescence of either alamarBlue cell viability reagent (Thermo Fisher Scientific) or Hoechst 33258 (Sigma-Aldrich) were measured by multi-detection microplate reader, POWERSCAN HT (BioTek). The quantification of melanin was carried out at OD475 with POWERSCAN HT. The total amount of melanin was normalized relative to the total viable cell number of the individual samples.

### siRNA treatment

ON-TARGET plus human CDH 1(E-cadherin) siRNA (Horizon Discovery) was used for E-cadherin knockdown. As a negative control, ON-TARGET plus Non-targeting Pool (Horizon Discovery) was used. siRNA treatment was performed according to the manufacturer’s instructions. Briefly, keratinocytes were transfected with 10 nM negative control or E-cadherin siRNA using Lipofectamine RNAiMAX (Thermo Fisher Scientific) in OptiMEM (Thermo fisher Scientific). After transfection, the cells were cultured for 24 or 48 hours and used on each assay. The siRNA-treated keratinocytes were further incubated by replacing the medium with the coculture medium (CnT-PRIME KM, CELLnTEC) for the coculture with melanocytes.

### Quantitative real-time PCR

Total RNAs were isolated from cells using the RNeasy Mini Kit (Qiagen) according to the manufacturer’s instructions. cDNAs were synthesized by using the PrimeScrip RT Reagent Kit (Perfect Real Time, TaKaRa). Quantitative real-time PCR (hereafter called qPCR) was performed by using the KAPA SYBR FAST qPCR Kit Master Mix (2×) Universal (KAPA Biosystems) with StepOne plus (Applied Bio systems). Primer sequences for qPCR were as follows: For human E-cadherin, forward (5’-CTGAAGTGACTCGTAACGAC-3’) and reverse (5’-ACGAGCAGAGAATCATAAGG-3’), for human Tyrosinase, forward (5’-TACGGCGTAATCCTGGAAACC-3’) and reverse (5’-CCGCTATCCCAGTAAGTGGA-3’), for human Tyrp1 forward (5’-GCTTTTCTCACATCGCACAG-3’) and reverse (5’-GGCTCTTGCAACATTTCCTG-3’), for human MITF, forward (5’-GCGCAAAAGAACTTGAAAAC-3’) and reverse (5’-CGTGGATGGAATAAGGGAAA-3’), for human GAPDH as reference gene, forward (5’-GGTGAAGGTCGGAGTCAACGGATTTGGTCG-3’) and reverse (5’-TATTGGAACATGTAAACCATGTAGTTGAGG-3’).

### Preparation of E-cadherin-knockdown conditioned medium

Keratinocytes were seeded at 4 × 10^4^ cells/well to a 12 well plate (Corning) and treated with either the control or E-cadherin specific siRNA for 24 hours. The medium was then replaced with CnT-PRIME KM containing 1 mM CaCl_2_ and incubated for 3 days. The supernatants of either the control or E-cad-KD keratinocyte culture were collected and centrifuged at 15,000 rpm for 5 min at 23 °C. For the mass spectrometry, the supernatants were specifically prepared as follows. After the knockdown, the medium was replaced with the coculture medium containing 1 mM CaCl_2_ and incubated for 24 hours. After incubation, keratinocytes were irradiated with or without 20 mJ/cm^2^ of UVB in the thermostatic UV irradiator (NK system) and incubated for another 24 hours. After 24 hours of incubation, the supernatants of the cultures (N=3/sample) were collected and centrifuged at 15,000 rpm for 10 min at 4 °C, and subjected to proteomic analysis. As an internal control, only the coculture medium was used to eliminate non-specific background proteins derived from the medium.

### Sample preparation for mass spectrometry

The EasyPep™ 96 MS Sample Prep Kit (Thermo Fisher Scientific) was used for sample preparation for mass spectrometry, including lysate preparation, protein digestion, and peptide enrichment and clean-up.

### LC-MS/MS analysis

The UltiMate 3000 RSLCnano system (Thermo Fisher Scientific) and Orbitrap Fusion Lumos (Thermo Fisher Scientific) were used as a LC-MS/MS system. A nano-HPLC capillary column (ODS, 75 μm i.d. x120 mm, particle size: 3.0 μm, Nikkyo Technos) was used. A linear 120 min gradient of 2−35 % mobile phase B was used for separation, where mobile phase A was 0.1% formic acid in water and mobile phase B was 0.1% formic acid in ACN. A data-dependent acquisition method using multiple filter criteria (charge state, monoisotopic m/z assignment and dynamic exclusion) for precursors was used. ITMS2 spectra were collected using a rapid scan rate with a maximum injection time of 50 ms. The LC-MS data was searched using Thermo Scientific™ Proteome Discoverer™ 2.5 software. By using the data from only the coculture medium and accessing identified peptide sequences together with derived species, protein compositions derived from the coculture medium were subtracted as the non-specific background.

### Paracrine effect on melanogenesis related factors

Melanocytes were seeded at 5 × 10^4^ cells/well to a 12 well plate (Corning) and cultured for 72 hours. The medium was replaced with either control or E-cad-KD conditioned medium. After 48 hours of incubation, RNAs were isolated and used for qPCR to examine the expression level of melanogenesis genes.

### Detection of melanin under coculture condition

Melanocytes were seeded at 2 × 10^4^ cells/well to a Φ27 glass-based 35 mm dish (IWAKI). At 24 hours after incubation, the melanocytes were cocultured with either control or E-cad-KD keratinocytes. After 7days of coculturing, the samples were fixed with 4% paraformaldehyde (PFA) in 1xPBS. The confocal laser scanning microscopy LSM880 (Zeiss) was used to observe melanin along with other fluorescent immunostainings.

### Time-lapse imaging of melanocytes

Melanocytes suspended in the medium were incubated with 0.5 mM Cell Tracker Green CMFDA Dye (Thermo Fisher Scientific) at 37°C for 25 min. The fluorescently labeled melanocytes were seeded to a Φ27 glass-based 35mm dish (IWAKI) and cultured for 24 hours. Keratinocytes were further cocultured with melanocytes so that the ratio of the total cell number between keratinocytes and melanocytes was 5:1. After 24 hours of coculturing, the culture dish was set in the stage top incubator (Tokai Hit, 37 °C, 5% CO_2_, 100% humidity) and time-lapse imaging was performed with a 2-min interval and 1 µm slice of z-stack for at least 6 hours by the confocal microscopy LSM880. Both the melanocyte cell-bodies and tips of dendrites were tracked using MTrack2 in ImageJ and the total migration trajectories were compared.

### Time-lapse imaging of melanin uptake

1% (w/v) of sonicated synthetic melanin (Sigma-Aldrich) in 1xPBS was added to keratinocytes at 24 hours after the knockdown treatment to the final condition of 0.002%. Immediately after the addition, time-lapse imaging was performed with a 15-min interval for 12 hours in a microscopic monitoring incubator BioStation CT (Nikon, 37 °C, 5% CO_2_, 100% humidity). The acquisition data was binarized using ImageJ.

### The screening of ingredients upregulating E-cadherin

Keratinocytes were seeded at 3 × 10^4^ cells/well to a 12 well plate and cultured for 48 hours. 0.005 % of individual ingredients diluted with DMSO were added to the cells. As a negative control, an equal amount of DMSO was added. After 24 hours of treatment, RNAs were isolated, followed by the quantification of E-cadherin mRNA expression by qPCR.

### Statistical analysis

The Data is presented as mean values ± SD. Statistical analyses were performed by either Student’s t-test (two-tailed) or one-way ANOVA with multiple comparisons. Differences with a value of *p* < 0.05 were considered statistically significant. For the proteomic analyses, all statistical analyses were performed using Perseus (version 2.0.3.1) [32].

### Immunofluorescent staining of cutaneous tissues and cultured cells

Fixed cultured cells and paraffin sections of cutaneous tissues were washed with PBST (1x PBS containing 0.1% Triton-X100) three times for 5 min each. The samples were further blocked with 2.5% goat serum in PBST for one hour at room temperature, and then incubated with appropriate primary antibodies (1:200 dilution for all primary antibodies in 1% goat serum with PBST) overnight. The following day, secondary antibodies for the appropriate species with the same dilution rate as in primary antibodies were incubated for 1.5 hours at room temperature. The nuclei were counterstained with DAPI (VECTASHIELD, Vector laboratories) for 10 min at room temperature. The static images of the immunostaining samples were retrieved with the 20x objective by the confocal microscopy LSM880, and further analyzed by ImageJ. The primary antibodies used were as follows: E-cadherin antibody (M361201-2, Dako), anti-Filaggrin (sc-66192, Santa Cruz Biotechnology), anti-Loricrin antibody (905104, BioLegend), anti-Trp1 antibody (TA99, Calbiochem), anti-ZO-1 antibody (33-9100, Invitrogen), anti-Desmoglein-1 (651110,Progen), anti-Desmocollin-1 antibody (HPA012891, ATLAS ANTIBODIES). The secondary antibodies used were as follows: Alexa Fluor 488 either anti-mouse or anti-rabbit IgG (A11001, A21206, Thermo Fisher Scientific), Alexa Fluor 647 either anti-mouse or anti-rabbit IgG (A21236, A21245, Thermo Fisher Scientific).

### Image processing and quantification of images

The data of the time-lapse imaging was analyzed by either image J or Imaris (Ver 8.4.1, BITPLANE). For the statistical analyses, all the images were binarized and measured by Image J.

## Results

### Downregulation of E-cadherin protein expression in age spots

Since dysfunctional differentiation of keratinocytes occurs in age spots [11][12], we first tested if cell adhesion molecules, which are important for epidermal differentiation are modulated in their expressions in age spots. By using cutaneous specimens of backs, we first examined the expression of an adherens junction molecule E-cadherin between non-lesion and lesion area (Fig. 1A). In the neighboring non-lesions, E-cadherin showed a clear epidermal expression in both basal and suprabasal layers (Fig. 1B). On the other hand, hyperpigmented lesions showed the decreased expression of E-cadherin in both basal and suprabasal layers (Fig. 1B, n=7). At the same time, the relative expression level of E-cadherin mRNA was also significantly decreased in age spots as compared with the non-lesions, suggesting that in age spots E-cadherin expression is downregulated at both mRNA and protein levels. To test whether E-cadherin downregulation is specific for age spots, other cell-cell interaction molecules were investigated by using age-spot specimens from faces (Fig.1A). For the homogeneous comparison, we compared fluorescent intensity between a perilesional and a lesional area on a same section. By comparison with the perilesional area, desmosomal cadherins Desmoglein-1(Dsg1) and Desmocollin-1(DSC1) showed no apparent change in their expressions in the lesion areas (Fig.1C, n=4). In addition, the tight junction molecule ZO-1 also showed no obvious difference between the perilesional and the lesion areas, indicating that the downregulation of E-cadherin is specific for age spots (Fig.1D, n=4). In this condition, terminal differentiation markers, Loricrin and Filaggrin were both decreased in their expressions in age spots, corroborating previous findings that differentiation is aberrant in age spots (Fig.1E, n=4).

**Figure 1.**
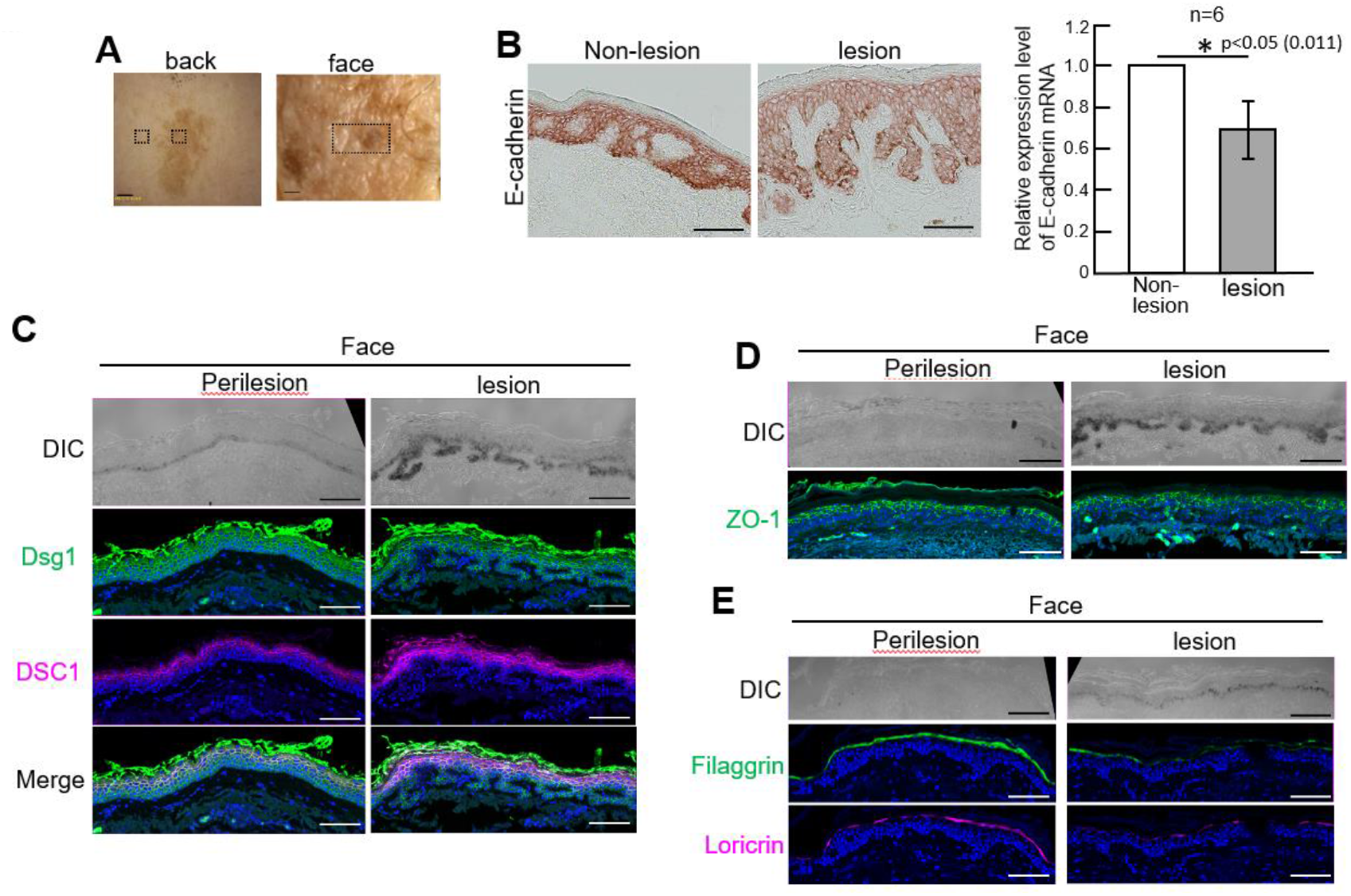
Decreased expression of E-cadherin in solar lentigines. **A**, Representative images of a solar lentigo (SL, back skin) and a hyperpigmented facial spot (see Materials and Methods). Dashed brackets denote examples of regions of interest analyzed in Fig.1B, C-E. **B**, E-cadherin proteins were decreased in the epidermis of solar lentigines (from the back skins). **C**, At mRNA level, E-cadherin was also decreased in solar lentigines. **D, E**, In comparison with neighboring non-lesion area, hyperpigmented lesion showed no obvious change in the expressions of other desmosomal cadherins, Dsg1 and DSC1 as well as a tight junction protein ZO-1. **E**, In comparison with the neighboring non-lesion area, the hyperpigmented lesion area showed decreased expressions of Filaggrin and Loricrin. DIC; Images of <u>D</u>ifferential Interference <u>C</u>ontrast. Scale bars, 1 mm (**A**), 100 µm (**B, C-E**). The data represents mean±SD.

Since E-cadherin is expressed in both keratinocytes and melanocytes, we next tried to distinguish an effect of E-cadherin downregulation on keratinocytes from its effect on melanocytes (Fig.2A). While E-cadherin downregulation by a neutralizing antibody against E-cadherin promoted the proliferation of keratinocytes as compared with control keratinocytes (Fig. 2A), the same treatment with the neutralizing antibody did not show any significant effects on melanocyte proliferation (Fig. 2B). In addition, E-cadherin neutralization had no effect on melanogenesis in either control or the neutralizing antibody-treated melanocytes (Fig. 2C). Therefore, these results suggested keratinocyte rather than melanocytes are considerably affected by E-cadherin inhibition. Intriguingly, a keratinocyte-melanocyte coculture treated with E-cadherin neutralizing antibody exhibited the promotion of melanogenesis with Trp1 upregulation (Fig. 2D), raising a possibility that melanogenesis are modulated through keratinocytes downregulated with E-cadherin.

**Figure 2.**
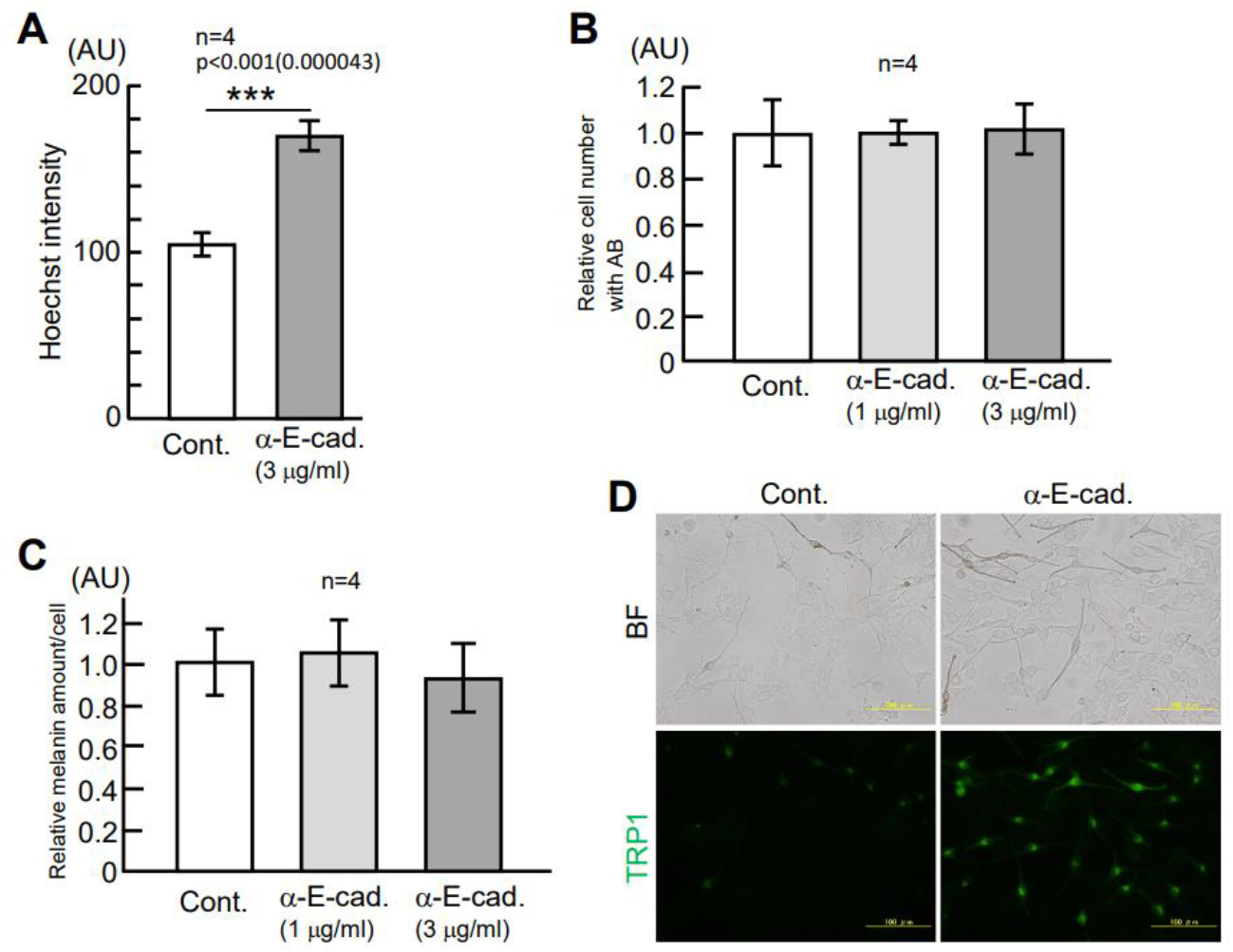
Promotion of keratinocyte proliferation and melanogenesis by neutralization of E-cadherin. **A, B**, Cultured keratinocytes, but not cultured melanocytes, significantly promoted proliferation in the inhibition of E-cadherin by a neutralizing antibody against E-cadherin (α-E-cad.). **C**, In the melanocyte monoculture, there was no effect on melanogenesis in E-cadherin neutralization. **D**, In the keratinocyte-melanocyte coculture, melanocytes promoted melanogenesis with upregulation of Trp1 protein expression, suggesting that the promotion of melanogenesis is dependent on the presence of E-cadherin-inhibited keratinocytes. BF; Bright field. Scale Bars, 100 µm. The data represents mean±SD.

### E-cad-KD keratinocytes promoted melanogenesis in a paracrine-dependent manner

Since neutralization of E-cadherin cannot necessarily determine a causal factor for the promotion of melanogenesis in the keratinocyte/melanocyte cocultures, we next performed specific knockdown of E-cadherin by siRNA only in keratinocytes. A siRNA against E-cadherin specifically depleted E-cadherin mRNA at the knockdown efficacy of over 96% in cultured keratinocytes (Fig. 3A). By using the coculture system, fluorescently-labeled normal melanocytes (Cell Tracker (MC)) were cocultured with either control or E-cadherin knockdown (E-cad-KD) keratinocytes, and external phenotypes of melanocytes were investigated. As in the above results with E-cadherin neutralizing antibody, melanocytes cocultured with E-cad-KD keratinocytes promoted melanogenesis with about 1.2 times more pigmented than that with control keratinocytes (Fig. 3B and 3C). On the other hand, melanocytes in both control and E-cad-KD keratinocytes’ cocultures showed no apparent changes in morphologies such as dendrite numbers, dendrite length, and the area of individual melanocytes (Fig.3D, 3E, and 3F). Therefore, these results indicated that melanogenesis is promoted by a manner dependent on the presence of E-cad-KD keratinocytes.

**Figure 3.**
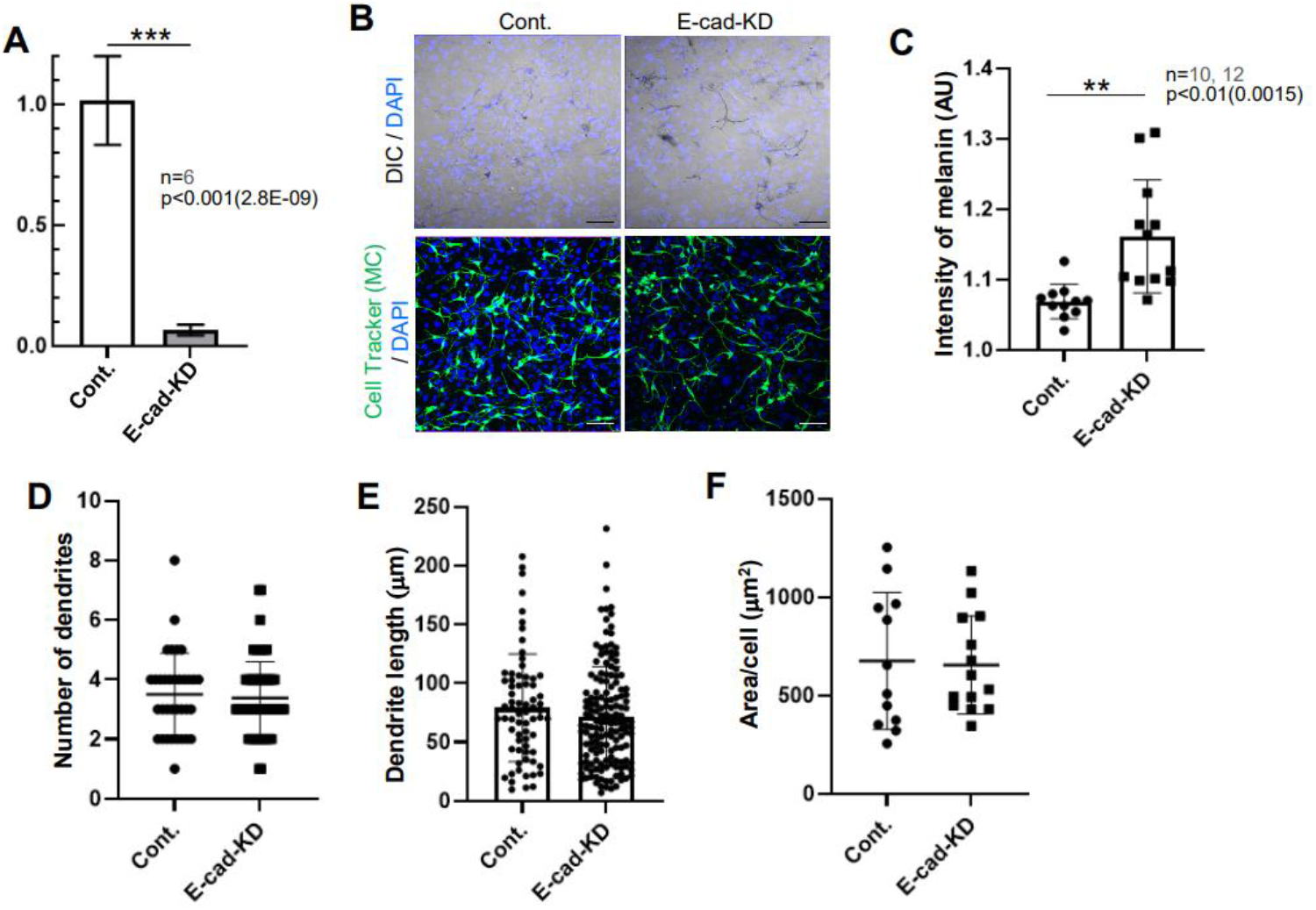
Melanocytes promote melanogenesis by coculture with E-cadherin knockdown keratinocytes. **A**-**C**, By coculture with either control or E-cadherin knockdown keratinocytes (E-cad-KD), melanocytes cocultured with E-cad-KD keratinocytes promoted melanogenesis more than ones cocultured with control keratinocytes. **D**-**F**, Melanocytes cocultured with both control and E-cad-KD keratinocytes showed no obvious morphological changes in the total number of dendrite per cell, the length of individual dendrites, and the area per cell. Scale Bars, 100 µm. The data represents mean±SD.

Since E-cadherin regulates cell-cell adhesion, there are two possibilities of the mechanism for promoting melanogenesis by E-cadherin downregulation: (1) weakening cell-cell interaction physically stimulates melanogenesis, (2) unknown secretion factors stimulate melanogenesis. To first test the latter possibility, we prepared a conditioned medium from the culture of keratinocytes treated with either control or E-cadherin siRNA (Fig. 4A, see Materials and Methods). Melanocytes were then cultured in the conditioned medium supplemented with 1 mM CaCl_2_ to moderately differentiate keratinocytes for activating E-cadherin, analyzed the mRNA level of melanogenesis genes. As a result, in melanocytes cultured in E-cad-KD conditioned medium, Tyrosinase and Trp1 mRNAs were significantly upregulated at about 1.2 times higher than those in melanocytes cultured in the control conditioned medium (Fig.4B). Also, MITF mRNA reproducibly tended to be upregulated via E-cad-KD conditioned medium (Fig.4B). Therefore, these results suggested that downregulation of E-cadherin triggers unknown secretory factors, which upregulate melanogenesis.

**Figure 4.**
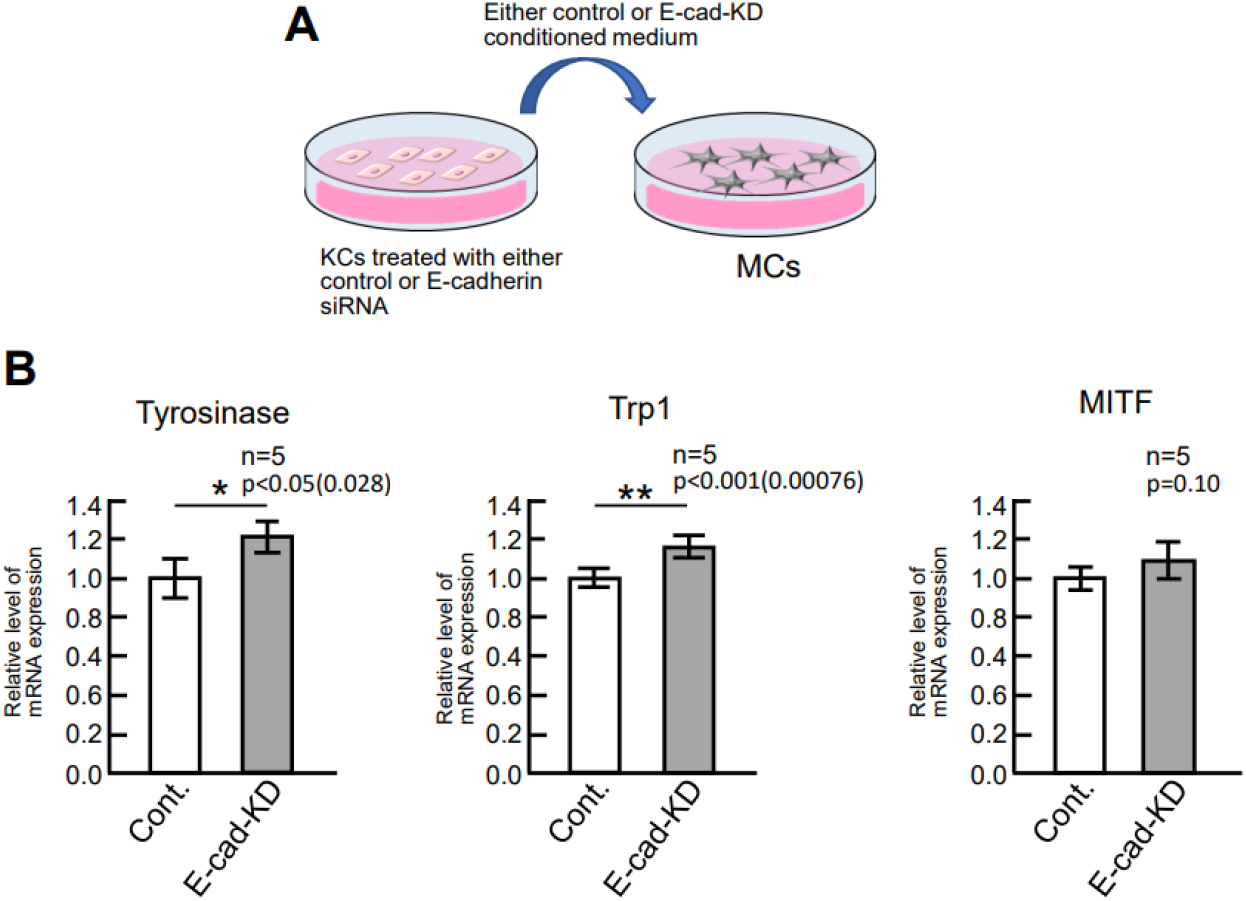
Upregulation of melanogenesis genes through paracrine factors of E-cad-KD keratinocytes. **A**, A schematic shows the melanocyte culture in either control or E-cad-KD conditioned medium. **B**, Tyrosinase and Trp1 mRNAs were significantly upregulated in E-cad-KD conditioned medium at about 1.2 times higher than those in the control. Also, MITF was reproducibly upregulated in E-cad-KD conditioned medium. These results suggested that unknown paracrine factors in E-cad-KD conditioned medium promotes melanogenesis. The data represents mean±SD.

### Downregulation of E-cadherin lead to the secretion of inflammatory and melanocytic factors

Melanogenesis is known to be stimulated by pleiotropic stimulating factors. Indeed, after UVB exposure, keratinocytes start secreting various melanocytic/inflammatory factors such as α-MSH, ET-1, IL-1α [14]. To investigate whether E-cadherin-KD triggers such secretory factors, we next performed proteomic analysis by using the supernatant of either control or E-cad siRNA-treated keratinocyte culture (the same as the above conditioned media) to identify secretory factors. To create a similar situation of the sun-exposed epidermis, siRNA-treated keratinocytes were irradiated with 20 mJ/cm2 UVB to stimulate downstream signaling molecules after UVB exposure. Principal component analysis and the heat map view of identified proteins showed clear difference in the identified protein composition between control and E-cad-KD conditioned medium (Fig. 5A and 5B). In addition, GO-term analysis showed that majority of proteins was secretory and exosome components (e.g., an exosome marker CD63 in Fig. 5D), demonstrating that secretory keratinocyte-derived factors were identified. By comparison with UVB-unexposed samples, UVB-exposed control-KD keratinocytes promoted the secretion of inflammatory factors such as TNF1α, MMP-9, and IL-1α (Fig.5C and D). In this condition, UVB-exposed E-cad-KD keratinocytes significantly upregulated both known inflammatory and melanocytic factors with abundance ratio ranging from over two to 20 times higher than that of UVB-exposed control conditioned medium (Fig. 5D). Among them, VEGFA, GM-CSF, and SCF are known to be involved in the process of pigmentation [5][6][22]. Since VEGFA, SCF, IL-1α, and SFRP1, are known to be upregulated in age spots ([22][33][34][35], Fig.5C and 5D), these results provided a key molecular insight into E-cadherin downregulation of keratinocytes that triggers various melanocytic and inflammatory factors for promoting melanogenesis as well as melanocytic environment such as chronic microinflammation.

**Figure 5.**
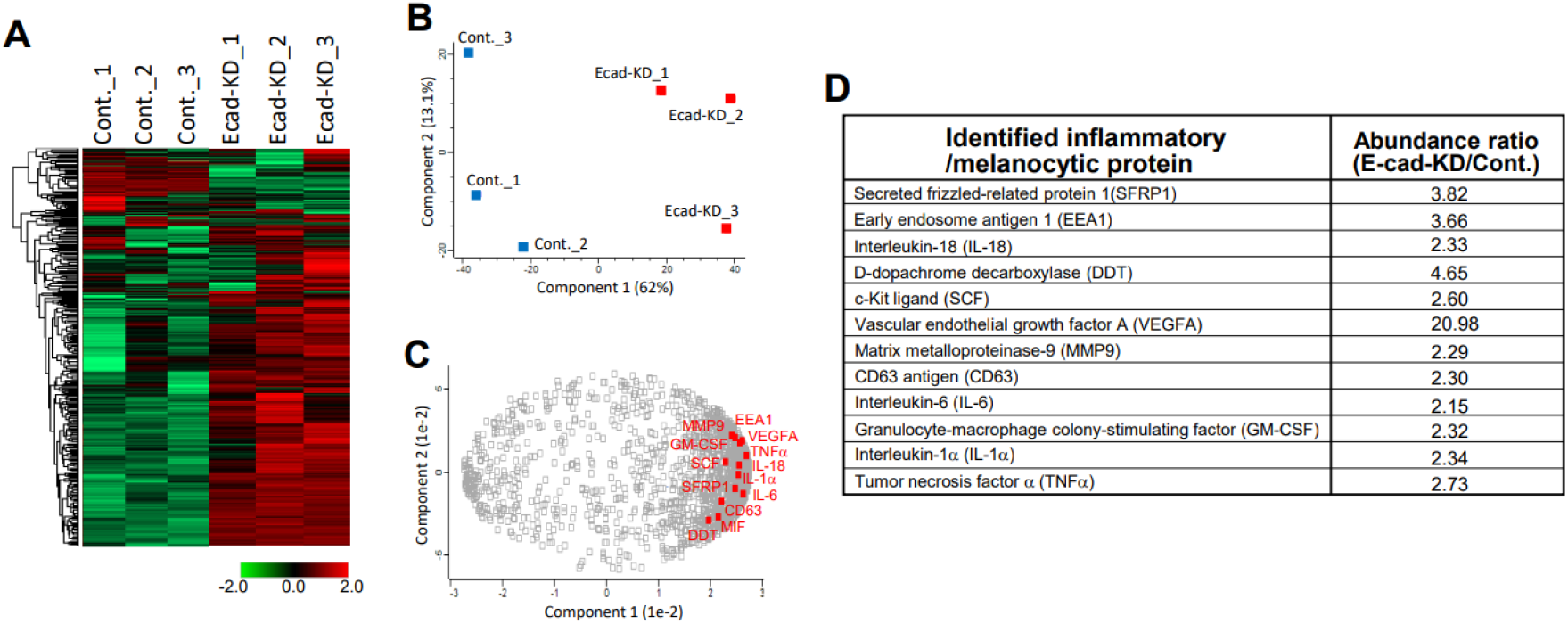
Identification of inflammatory and melanocytic factors upregulated by E-cadherin downregulation. **A**, A heatmap view of identified proteins (2706 proteins identified) in the supernatant of either control or E-cad-KD keratinocytes by mass spectrometry (n=3 each). **B**, Principal component analysis shows clear difference between control and E-cad-KD supernatants. **C**, Identified inflammatory and melanocytic proteins were strongly related with the components of identified proteins in E-cad-KD supernatants. **D**, In E-cad-KD supernatants, known inflammatory and melanocytic factors involved in age spots and pigmentation were identified and upregulated.

### Stagnant dynamics of melanocytes in the presence of E-cad-KD keratinocytes

Although the paracrine effect of E-cadherin downregulation on melanocytes were revealed, how the cell adhesive function of E-cadherin affects on melanocytes has remained to be addressed. Since E-cadherin plays a crucial role in cell migration via its function of cell-cell contact [31][36], we next tested whether downregulation of E-cadherin also affects the behavior of melanocytes such as cell migration and dendrite outgrowth. To this end, live imaging of melanocytes cocultured with keratinocytes were carried out. While control melanocytes appeared to actively migrate with active dendrite outgrowth and retraction, the melanocytes cocultured with E-cad-KD keratinocytes less migrated with rather static dendrite outgrowth (Fig. 6A and 6B, the top panels). In fact, the speed of both melanocyte migration and the retraction/growth rate of dendrites in E-cad-KD coculture were significantly slower than those of the control based on the distance of either cell body or dendrite trajectory (Fig. 6A and 6B, the bottom graphs). Therefore, these results indicated that cell-cell contact mediated by E-cadherin is important for a proper melanocyte-dynamics. The stagnation of melanocyte dynamics may generate a higher probability of timing for transferring melanin to keratinocytes.

**Figure 6.**
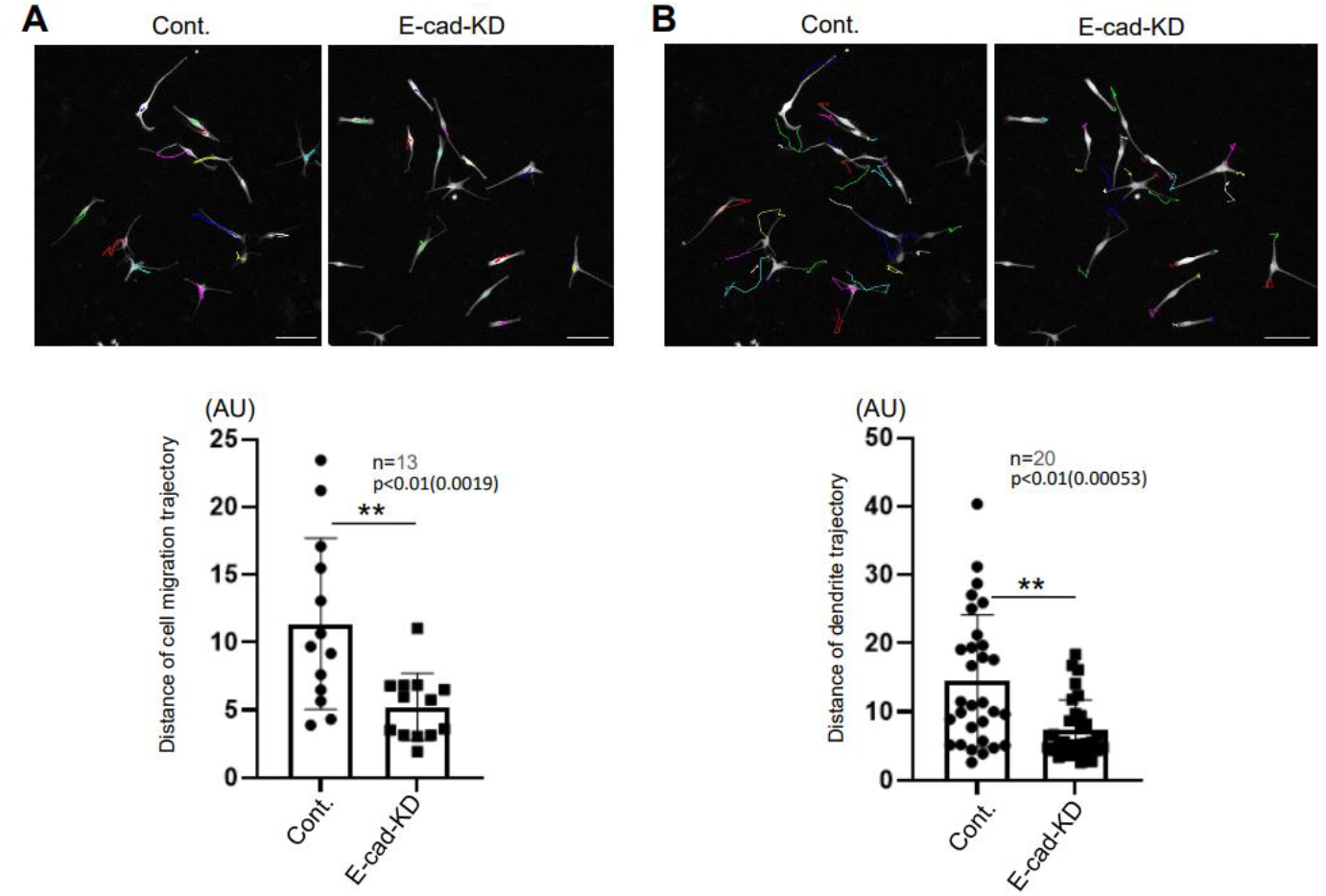
Stagnant melanocyte dynamics in coculture with E-cad-KD keratinocytes. **A, B**, By tracking the cell body and the tips of dendrites of individual melanocytes, cell migration and dendrite retraction/outgrowth were both significantly prevented in the presence of E-cad-KD keratinocytes but not control ones. Scale Bars, 100 µm. The data represents mean±SD.

### Promotion of melanin uptake by E-cadherin downregulation

E-cadherin downregulation prevented normal keratinocyte differentiation, thereby maintaining proliferative state of keratinocytes (Fig. 2A, [28]). Since proliferative basal keratinocytes more actively ingest and store melanin than differentiated keratinocytes [37], we further investigated the speed of melanin uptake of either control or E-cad-KD keratinocytes. Immediately after adding 0.002% of melanin, time-lapse imaging of keratinocytes under bright field microscopy were performed. After 2h of incubation, melanin uptake started, and the rate of the melanin uptake maximally became faster than that of the control (Fig. 7A and 7B). The maximum rate of change in the speed of melanin-uptake in E-cad-KD keratinocytes was about two times faster than that in control keratinocytes (Fig. 7B). Therefore, these results suggested that by downregulation of E-cadherin undifferentiated-keratinocytes prone to promote melanin uptake, which may also contribute to constitutively form melanin-stored keratinocytes in age spots.

**Figure 7.**
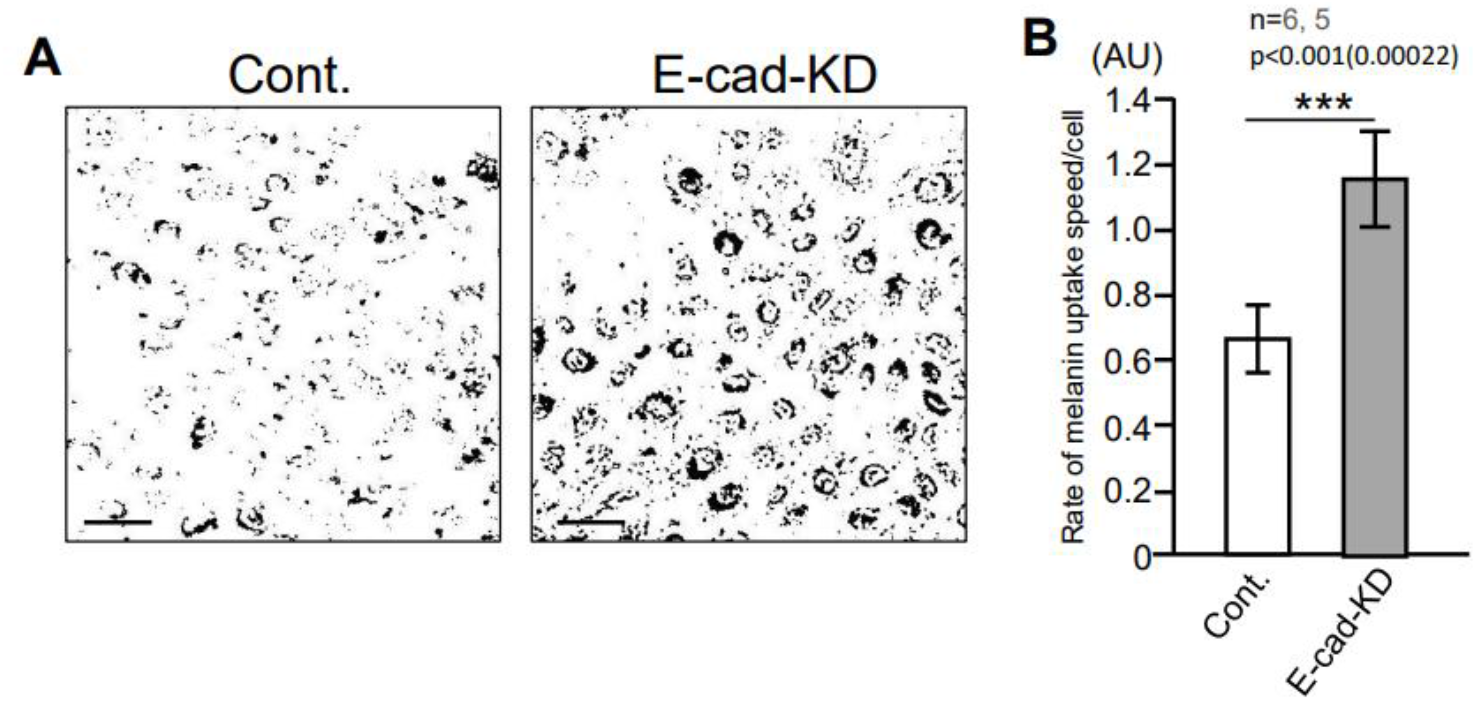
Promotion of melanin uptake in E-cad-KD keratinocytes. **A**, A binarized bright field image of either control or E-cad-KD keratinocytes shows the accumulation of melanin around nuclei at 2h after melanin addition. **B**, After 2h of melanin addition, E-cad-KD keratinocytes significantly promoted the rate of melanin-uptake speed at about 1.5 times faster than that of control keratinocytes. Scale Bars, 50 µm. The data represents mean±SD.

### A solution for upregulating E-cadherin by Rosa multiflora fruit extract

E-cadherin downregulation in keratinocytes was involved in multiple melanocytic processes (see also Fig. 8B): (1) the promotion of secreting inflammatory and melanocytic factors, (2) the promotion of melanogenesis, (3) the modulation of melanocytes dynamics, (4) the promotion of melanin uptake. Given that E-cadherin was specifically decreased in age spots, it is plausible that prevention of these processes all together could be an integrated solution to improve age spots. One simple solution is upregulation of E-cadherin by plant extracts or chemical ingredients. Therefore, to provide such solutions, we finally carried out the screening of ingredients. By treating cultured keratinocytes with various percentage of ingredients, keratinocytes were subjected to quantitative PCR. As a result, we found that 0.005 % of Rosa multiflora fruit extract was able to upregulate E-cadherin mRNA at 1.2 times higher than the control (Fig.8A). Thus, this result provided a first example for the solution that could approach the origin of downstream processes of age spots, E-cadherin downregulation.

**Figure 8.**
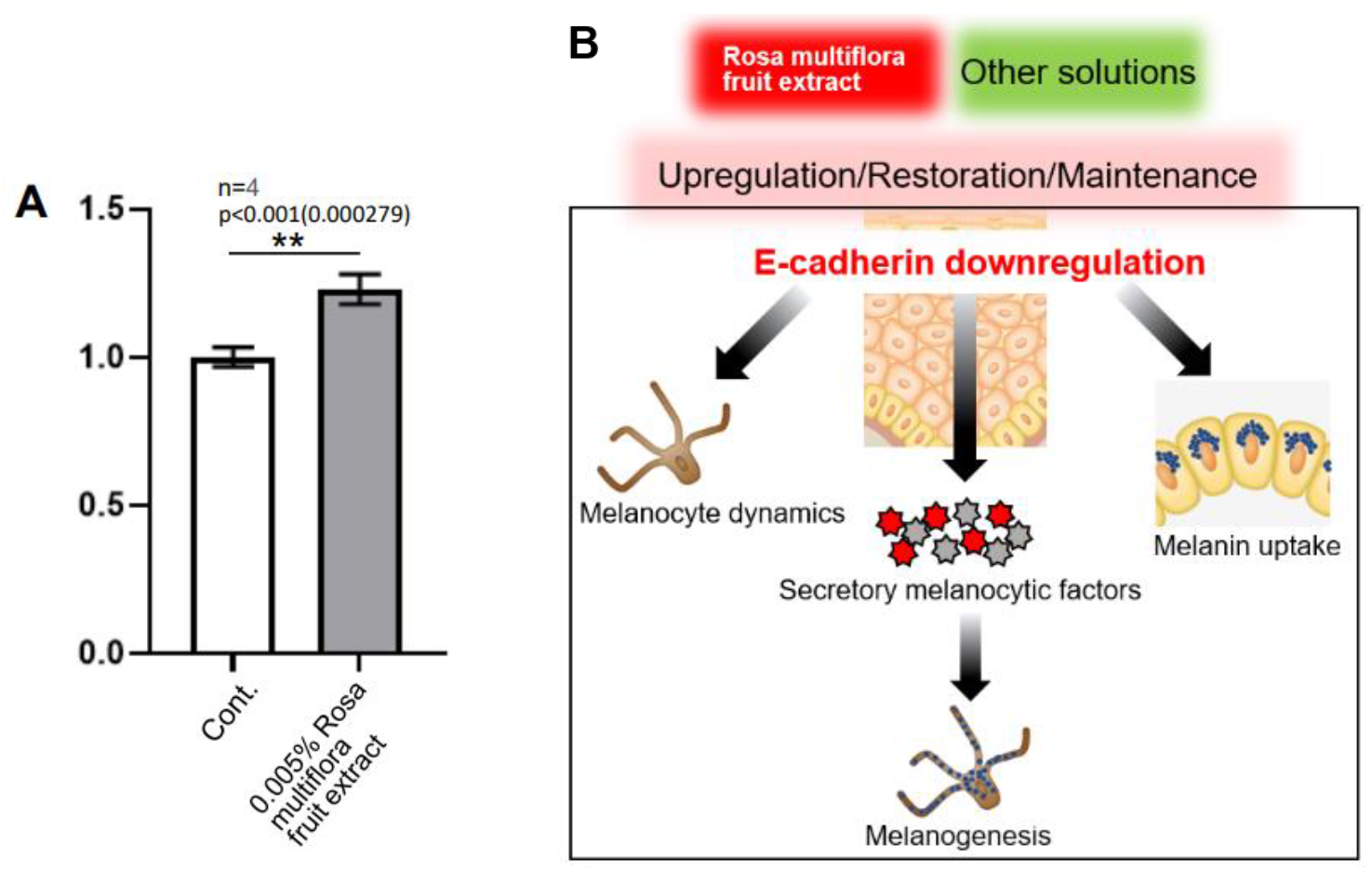
Upregulation of E-cadherin by Rosa multiflora fruit extract and a perspective based on the findings. **A**, By a screening for ingredients upregulating E-cadherin, 0.005% Rosa multiflora fruit extract was able to upregulate E-cadherin mRNA at about 1.2 times higher than the control vehicle. The data represents mean±SD. **B**, A schematic representing the core findings in the research. E-cadherin downregulation in age spots could trigger four downstream melanocytic processes in both melanocytes and keratinocytes. Besides Rosa multiflora fruits extract, future solutions could approach not only upregulating E-cadherin, but also restoring and maintaining E-cadherin to prevent the melanocytic origin.

## Discussion

Long-standing research for age spots has revealed the age-spot specific mechanisms and solutions. So far, these studies are categorized as below molecular processes: (1) Failure of keratinocyte differentiation [11][12], (2) Upregulation and secretion of various inflammatory and melanocytic factors [6][13][14], (3) Promotion of melanogenesis by melanocytic factors [15][16], (4) Degradation of basement membrane by inflammatory factors [17][18]. Previous studies have investigated the above individual processes for age spots, showing respective responsible factors. For example, the inhibition of tyrosinase, is one of the major ingredient solutions for age spots, but only targeting melanocytes [38][39]. Besides melanocytes, surrounding cells such as keratinocytes and fibroblasts were also the targets for solutions, which are directly or indirectly interacted with melanocytes for melanogenesis. Particularly, after UVB exposure, secretory factors from keratinocytes have been also targets for cosmetic ingredients [38][39]. Recently, new approaches with optical coherence tomography (OCT) and clearing tissue technique found microstructure underlying age spots such as hyperplasia of micro-vascularization and cutaneous nerve fibers, demonstrating that the interaction between other tissue systems and cutaneous system is critical for understanding basic mechanism of age spot formation and its maintenance [23][40][41]. However, how individual processes are related or integrated with each other has remained elusive. Our present study revealed that downregulation of E-cadherin could be an upstream cause for forming chronic melanocytic environment (Fig.8B). E-cadherin downregulation in keratinocytes could first start upregulating the secretion of inflammatory and melanocytic factors from keratinocytes. These secretory factors then stimulate melanocytes for promoting melanogenesis. Furthermore, probably due to unstable cell-cell contact, melanocytes less migrate with rather static elongated dendrites. Due to its stagnant state, melanocytes may have more chance to transfer melanin to keratinocytes rather than normal ones. In addition to this, we further showed that proliferative E-cad-KD keratinocytes were prone to promote melanin uptake. Thus, downregulation of E-cadherin triggers various downstream melanocytic processes, secretion of melanocytic factors, forming a “loop” between keratinocytes and melanocytes. In other words, E-cadherin is a “fort” of keratinocytes that prevents epidermis from triggering melanocytic factors as well as forming the inflammatory environment.

Accumulating evidence has shown that E-cadherin downregulation is involved in melanoma metastasis and vitiligo [31] [42][43]. For example, E-cadherin downregulation in melanocytes leads to melanocyte detachment from the basement membrane, leading to the apoptosis of melanocytes [42][43]. On the other hand, melanoma alternately expresses N-cadherin by losing E-cadherin, thereby infiltrating into the dermis layer [44]. Therefore, E-cadherin in other cell-types is also the critical causal factor for cutaneous disorders. The same is true in keratinocytes, severe ionizing radiation caused the decreased expression of E-cadherin protein, leading to radiation dermatitis [45]. At the daily life level, UVB and mechanical stresses could accompany reorganization of cell-cell contact in the epidermis via the cleavage or internalization of E-cadherin [46][47]. Therefore, cutaneous homeostasis is elaborately regulated by E-cadherin, downregulation of which could lead to various cutaneous abnormalities. Considering age spots and vitiligo, E-cadherin seems to be a double-edged molecule for hyperpigmentation and hypopigmentation [42][43]. Since our study showed that E-cadherin downregulation in keratinocytes triggered the secretion of many inflammatory and melanocytic factors, it is possible that in vitiligo melanocytes and surrounding keratinocytes secret a different composition of secretory factors. Therefore, for improving both macular disorders, it is crucial to pin down detailed compositions of secretory factors from both E-cadherin-downregulated melanocytes and keratinocytes.

While intensive studies have uncovered various factors responsible for age spot formation, how those factors reciprocally are related or integrated yet addressed. Thus, to improve age spots, the integrated solution targeting the origin of age spot has been sought out. By revealing E-cadherin as a novel core factor of age spots, our finding opens a new possibility for developing an integrated solution. As a first solution, we showed that Rosa multiflora fruit extract increases E-cadherin expression in keratinocytes. Strengthening cell-cell contact by restoring E-cadherin expression could be the integrated solution that could cover various downstream melanocytic-environments. Regarding restoration of E-cadherin expression, many natural ingredients such as Melatonin, Metformin, and Resveratrol are considered for cancer therapy by restoring E-cadherin expression along with inhibiting malignancy factors [48][49]. On the other hand, the prevention of E-cadherin downregulation mediated by UVB and mechanical stresses could also be another aspect of the solution [46][47]. Therefore, both restoration and maintenance of E-cadherin expression could be pragmatic solutions for not only improving age spots but also preventing the potential risk of the age spot formation (Fig. 8B).

## Conclusion

We showed that E-cadherin downregulation of keratinocytes in age spots could be the origin of key processes for the chronic melanocytic environment: upregulation of secretory melanocytic/inflammatory factors, upregulation of melanogenesis, upregulation of melanin uptake, and detention of melanocyte dynamics. Based on the hub function of E-cadherin, the first inclusive solution to improve age spots, Rosa multiflora fruit extract was provided.

## Acknowledgments

We are grateful to the emeritus professor Hachiro Tagami, Tohoku University, as dermatology specialist for assessing solar lentigo and providing insightful comments for the research. We also thank colleagues of MIRAI technology institute, Shiseido Co. Ltd. for the helpful discussion.

